# Adaptive evolution of phenotypic switching during environmental change

**DOI:** 10.1101/2020.06.02.127241

**Authors:** Urvish Trivedi, Maria R. Rebsdorf, Angel R. Cueva, Kendra P. Rumbaugh, Mette Burmølle, Søren J. Sørensen, Jonas S. Madsen

## Abstract

Microbes are faced with environmental fluctuations and it is therefore important to understand how change affects bacterial evolution and population dynamics. Here, we evolved a model bacterium, *Pseudomonas aeruginosa* PA14, under both constant and changing conditions where settlement by biofilm formation or migration by swimming motility was induced. In changing environments, a heterogeneous population evolved with both specialists and generalists. Interestingly, over time, generalists were outcompeted despite being better at both phenotypes compared to the ancestor. However, generalists became dominant when faced with local competition after resettlement. Reintroduction into a burn wound model, the environment PA14 originally was isolated from, further suggested a tradeoff between niche breadth and peak fitness in agreement with the evolution experiments. Mutations in the c-di-GMP system were key for adaptation of the phenotypes. No mutants lost their ability to regulate c-di-GMP, but generalists acquired mutations that optimized their phenotypic response by efficiently adjusting c-di-GMP levels.

## Introduction

Evolution of organisms is closely tied to the level of stability in their local environment. Most environments are dynamic and generalist organisms have therefore evolved to sense and respond to environmental change by switching their phenotype and thereby broadening their niche space. Responsive phenotypic switching can evolve if the environmental change is recurring, as the organism typically needs to sense the environmental fluctuation to express the phenotype that makes it most fit^1^. Alternatively, niche differentiation may occur as a response to environmental change. Here, the population differentiates into co-existing sub-populations that specialize and thus optimize their fitness to different sub-niches. However, this type of specialization may make the organism ill-adapted in alternative niches. Niche specialization, therefore, is typical in stable environments, where such a tradeoff has little effect^2–4^. Thus, the type of phenotype and environmental change affects what evolutionary strategy facilitates optimal fitness tradeoffs^5^. Here the aim was to study adaptation in changing environments by experimental evolution using bacteria, as this approach is notably underutilized within this important research field. Experimental evolution enables empirically-based and direct testing of theory by using organisms with short generation times, that can be cultivated in highly controlled environments, thus enabling adaptation to occur and be examined in real-time.

Here a central responsive phenotypic switch of bacteria was studied, namely the transition between a sessile (e.g. biofilm) and a planktonic state (e.g. swimming). Biofilms are sessile microbial communities encased in an extracellular polymeric matrix. Most bacteria are able to shift from a sessile biofilm state to a planktonic state, where the bacteria typically become singular and motile if proficient. Over the last two decades, research has focused on understanding the biofilm state and the molecular adaptations that occur upon biofilm formation^6–8^, whereas the knowledge about adaptations to the planktonic motile state is more limited^9^. In addition, very little is known about how bacteria adapt towards responsive switching itself and thus between the two states. Therefore, here we studied the adaptation of this responsive phenotypic switch using the model organism *Pseudomonas aeruginosa* PA14.

We performed experimental evolutions where we alternated between biofilm formation and swimming motility: Static broth cultures were used to induce biofilm formation as *P. aeruginosa* PA14 forms thick biofilm mats, termed pellicles, at the air-liquid interface under this condition. Previous studies have shown that pellicle formation enables the bacteria to access oxygen, the energetically most favorable electron acceptor *P. aeruginosa* can utilize. Pellicle specialists can, therefore, outcompete non-specialists in static broth cultures^4,10^. Swimming motility was induced in low-nutrient soft agar medium. Here the actively metabolizing bacteria establish a nutrient gradient and swimming motility is triggered via chemotaxis. The further a bacterium can swim, the more access it has to nutrients, which is the basis of the selective pressure in the swimming assay^11^. The same basic nutrients were available in both assays.

## Results

### Stable environments select for specialists with fitness tradeoffs in opposing phenotypes

First, control experiments that mimicked stable environments – bacteria propagated in only biofilm or swimming assay – were examined. After a specified number of passes, the colony morphology and swimming phenotype of between 30 to 60 random isolates was tested. The colony morphology was assessed using a Congo red based assay, where a more wrinkled and red colony morphology is associated with elevated expression of the Pel matrix component and therefore correlated with increased biofilm formation^4^. Isolates with similar colony morphology and swimming phenotype (diameter) were assumed to be the same cell lineage (genotype). Figure 1a (Fig. S1) illustrates how, compared to the ancestor (*A^0^*), the isolates from the biofilm (*C^B2^*) and swimming (*C^S1^*) control experiments had a more pronounced phenotype; more wrinkled and redder colony morphology or larger swimming diameter, respectively. The isolates from these control experiments that were not selected for had consistently less pronounced phenotypes, suggesting a fitness tradeoff between the two phenotypes in question, which is in agreement with the physiological nature of the two phenotypes. Indicating that the biofilm and swimming control experiments selected for specialists.

**Figure 1.**
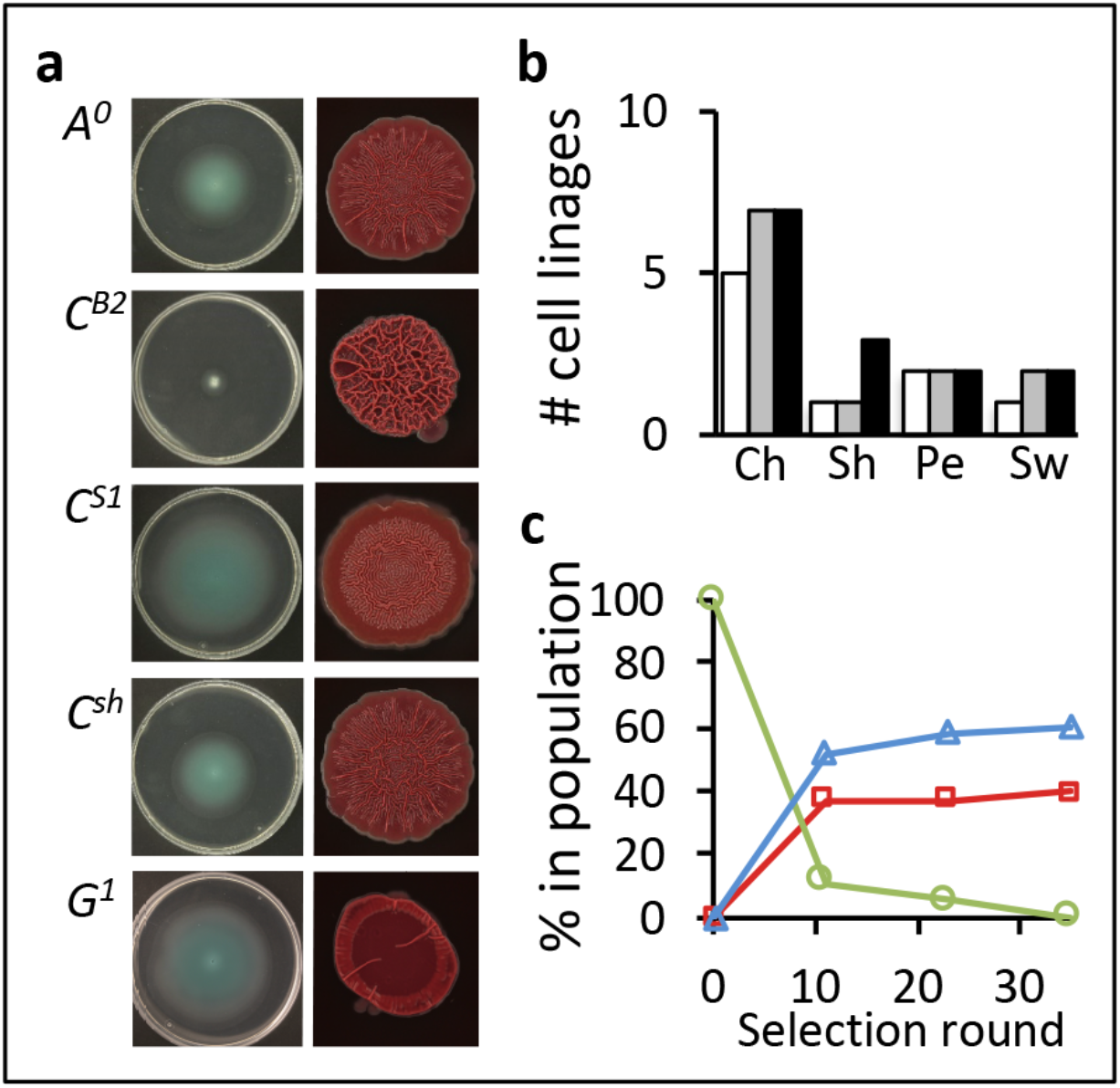
Environmental change facilitates phenotypic heterogeneity where generalists were outcompeted over time. **a**) Examples of colony morphology and swimming motility of cell lineages. *A^0^*, ancestral PA14; *C^B2^*, biofilm specialist from control experiments in the pellicle assay; *C^S1^*, swimming specialist from control experiments in the swimming assay; *C^sh^*, generalist from control experiments in shaken cultures; *G^1^*, generalist from experimental evolution in changing environments. **b**) Number of different cell lineages observed in changing environments (Ch), shaken cultures (Sh), the pellicle assay (Pe), and swimming assay (Sw). White bars correspond to selection round 11, gray bars selection round 23, and black bars selections round 35. **c**) The proportion of generalists (green), biofilm specialists (red), and swimming specialists (blue) observed during experimental evolutions in changing environments over time.

An additional control experiment was performed, where bacteria were propagated only in shaken cultures. Here, the vast majority of isolates resembled the ancestral strain, confirming that the biofilm and swimming assays selected for the phenotypes in question (Fig. 1a; *C^sh^*) whereas shaken cultures did not.

### Changing environments select for heterogeneity

When examining isolates from experiments where bacteria were passed between the biofilm and swimming assays, mimicking changing environments, different lineages emerged. Compared to the control experiment, it was clear that environmental change selects for a more heterogeneous population (Fig. 1b). After changing the environment 11 times (e.g. 6 times in both the swimming and biofilm assay) both generalists and specialists were found; however, during selection for a longer period (35 rounds), the generalists were outcompeted (Fig. 1c). To further characterize generalists and specialists, the competitive fitness of representative isolates of the distinct lineages was tested, relative to the ancestral strain, both in the biofilm and the swimming assay (Fig. 2, Table S1a). The competitive fitness of the generalists (*G^1^* and *G^2^*) was enhanced compared to the ancestor in both the biofilm and swimming assay. Comparatively, the specialists had higher fitness in the assay that favored their phenotype but lower fitness in the opposing assay.

**Figure 2.**
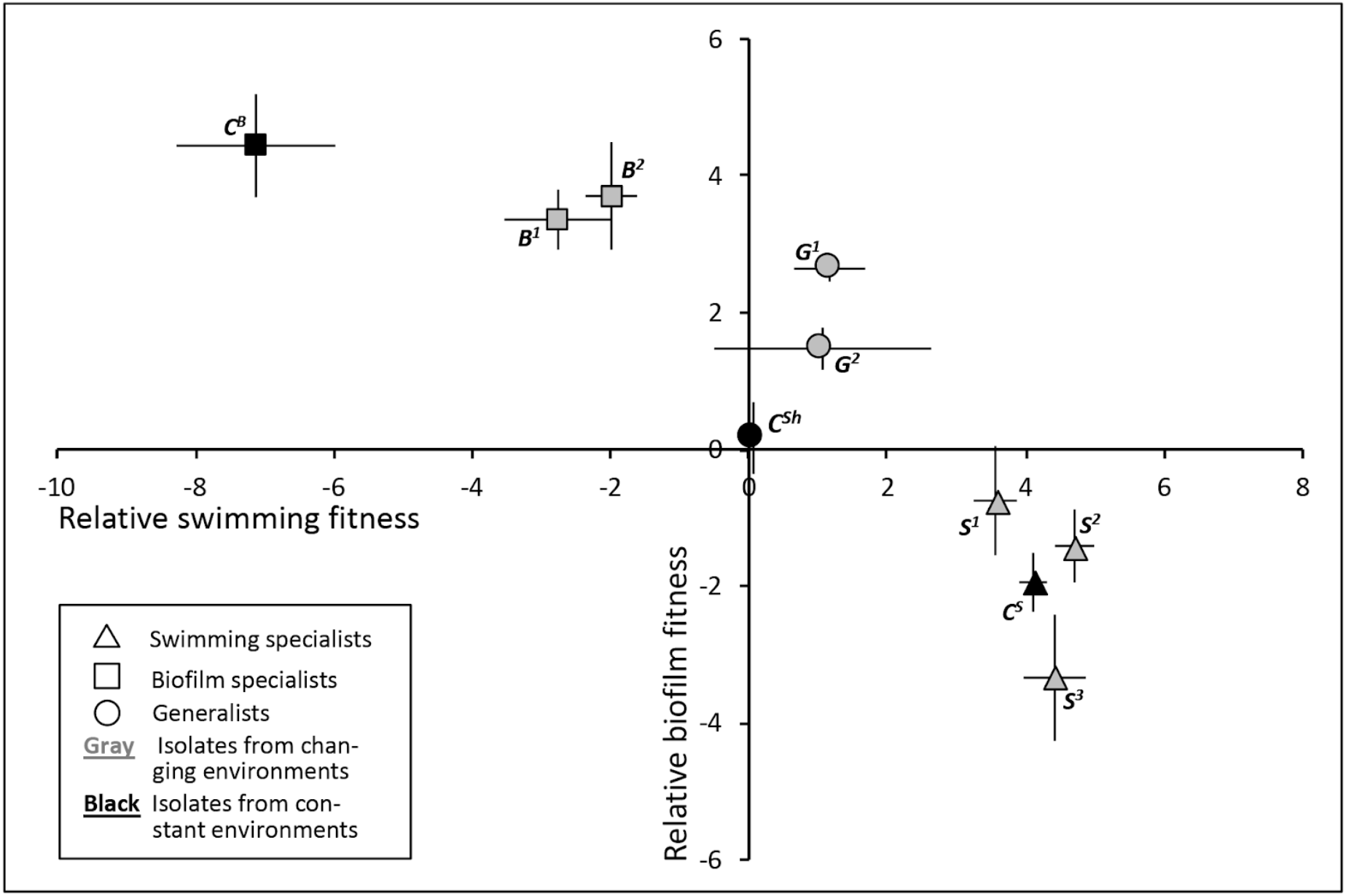
Generalists with enhanced biofilm and swimming phenotypes were observed and fitness tradeoffs occurred among specialists. The competitive fitness of representative isolates of the unique lineages measured in the swimming and the biofilm assay is shown relative to the ancestral strain *A^0^*. Data is based on 6 independent replicates and was Log_2_ transformed. Error bars represent the standard deviation. See Table S1a for *P*-values of one-way ANOVA with post hoc Dunnett test.

### Early molecular adaptation of specialists and generalists

The genomes of one representative isolate from each of the unique lineages from the evolution experiment initiated with the original ancestor were genome sequenced to characterize the mutational adaptation of the strains (Table S2). Those from the changing environments had a mutation in *fleN* (FleN_175c>G_). This specific mutation leads to multi-flagellation and has previously been shown to increase swarming and swimming motility of *P. aeruginosa* PA14^12^. The *fleN* mutation was the only one identified in swimming specialist *S^1^*.

Swimming specialists *S^2^* had, in addition to the *fleN* mutation, a mutation in the ribosomal binding site upstream of *groES*. This mutation further increased the swimming ability of *S^2^* compared to *S^1^* but also made *S^2^* less fit in the biofilm assay compared to the ancestor (Fig. 2, Table S1a). Missense mutations in swimming specialist *S^3^* included the two aforementioned mutations in addition to a deletion mutation in *PAI4_29800* (PA14_29800_262ΔRAG_), which encodes a methyl-accepting chemotaxis protein (28% similar to WspA). The sensory module for signal transduction of PA14_29800 is different from the one WspA holds. Biofilm specialist *B^1^* had, in addition to the *fleN* mutation, a mutation in *wspA* (Wsp_A382v>A_), a chemotaxis transducer of the Wsp system. Biofilm specialist *B^2^* had, besides *fleN* and *wspA*, mutated in a *lasR*-like gene PA14_45960 (PA14_45960_231A>T_). Mutations in generalists *G^1^* and *G^2^* were, in addition to that in *fleN*, identified in *wspF* (WspF_185Q>R_) and *wspA* (WspA_449F>c_), respectively.

Based on the identified mutations of the isolates of the experimental evolutions in changing environments, two transitions seemed important for early adaptation of the generalist with higher competitive fitness than the original ancestor, namely: 1) The *fleN* (FleN_175c>G_) mutation enabled multi-flagellation, not only in swimming specialists but also in biofilm specialists (Fig. S2). Multi-flagellation propels the cells further, compared to the ancestor with a single flagellum, when motility was induced. 2) Mutations in the Wsp chemosensory system, which are likely to affect the intracellular levels of c-di-GMP and hereby modify both swimming motility and the level of biofilm formation.

### Multi-flagellation and modified cyclic di-GMP levels enable specialization and enhanced responsive phenotypic switching of generalists

In accordance with our results, the specific mutation in *fleN* (FleN_175c>G_) has previously been shown to confer a reduction in biofilm formation (Fig. 2; *S^1^*)^12^. We, therefore, speculated that the Wsp mutations in strain *G^1^* and *G^2^* compensated for this by amending the intracellular levels of c-di-GMP. Levels of c-di-GMP were therefore gauged by introducing a promoter-GFP fusion based c-di-GMP reporter into the different mutants. Figure 3 (Table S1b) shows the level of c-di-GMP of each strain relative to the ancestor (*A^0^*) as a function of their competitive fitness against *A^0^* in the swimming and biofilm assays.

**Figure 3.**
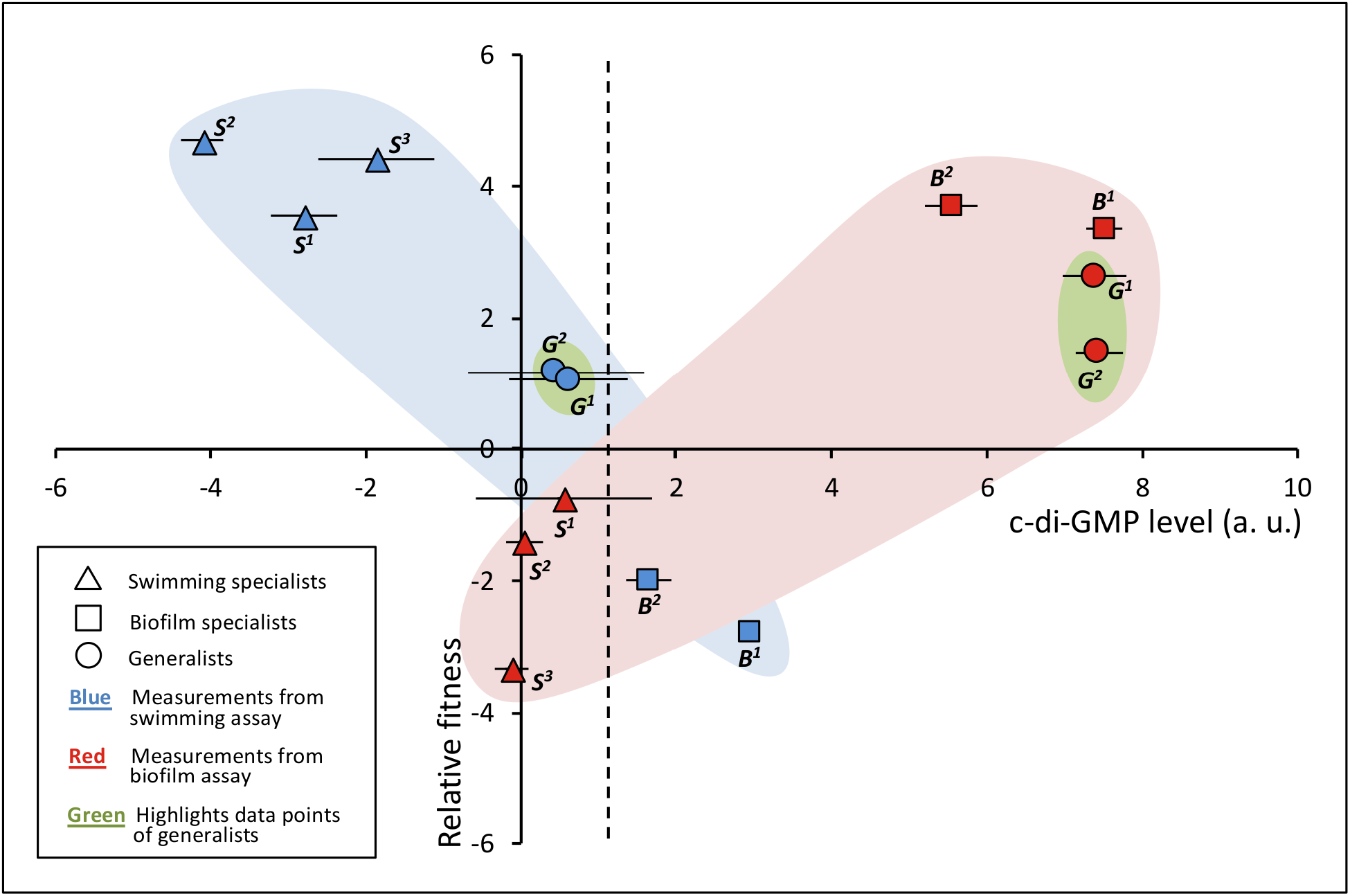
Cyclic di-GMP levels are interrelated with the competitive fitness of strains in the swimming and biofilm assay. The competitive fitness of isolates as a function of c-di-GMP levels, both relative to the ancestral strain *A^0^*. The stippled vertical line indicates where higher or lower level of c-di-GMP comes with fitness tradeoffs in the swimming and the biofilm assay, respectively. Cyclic di-GMP data is based on 4 independent replicates and was Log2 transformed. Error bars represent the standard deviation. See Table S1b for *P*-values of one-way ANOVA with post hoc Dunnett test.

Levels of c-di-GMP positively correlated with the competitive fitness in the biofilm assay (Fig. 3, **Pearson’s *r=* 0.85,** *P* = 0.007) and negatively correlated in the swimming assay (Fig. 3, **Pearson’s *r=* 0.94,** *P* < 0.001) revealing that the competitive fitness of the different strains was interrelated with levels of c-di-GMP. The c-di-GMP levels of biofilm specialists were considerably higher in the biofilm assay compared to the swimming specialist and the ancestor, however, their levels were also higher in the swimming assay. This indicates that the poor swimming ability of biofilm specialists, the fitness tradeoff, was due to a lack of proficiency in reducing c-di-GMP levels enough to fully switch to the planktonic swimming state. In accordance, the opposite appeared to be the case for swimming specialists whose c-di-GMP levels were lower than those of the biofilm specialists and generalists in the swimming assay. It appeared that the c-di-GMP levels of swimming specialists were roughly the same as the ancestor in the biofilm assay, despite their lower fitness. Interestingly, the levels of c-di-GMP were also higher among generalist *G^1^* and *G^2^* in the biofilm assay, matching those of the biofilm specialists. However, *G^1^* and *G^2^* were able to reduce c-di-GMP levels to below those of the biofilm specialists (Fig 3; stippled vertical line), roughly to the same levels as the ancestor. The range from low c-di-GMP levels in the swimming motility assay and the high levels in the biofilm assay was largest among the generalist, which indicates that, subsequent to becoming multi-flagellated, the generalists adapted by better adjusting their c-di-GMP levels in response to the environment.

### Selection for generalists - resettlement, spatial isolation and local competition

The above suggested that a generalist strategy was not viable during global competition against both biofilm and swimming specialists. As motility is one of the phenotypes of interest, dispersal, and resettlement are relevant ecological factors to consider: Motility and other planktonic states enable bacteria to disperse and resettle in new environments and hereby become spatially isolated from competing lineages.

Therefore, we mimicked resettlement by restarting the experimental evolutions in changing environments, beginning with the adapted generalists (*G^1^* and *G^2^*) as the ancestors. The colony morphology and swimming performance of isolates were screened after 11 passes, corresponding to the first time point of the experimental evolutions started with the original ancestor (Fig. 1c). We found that the diversity was as high as in experiments initiated with the original *P. aeruginosa* ancestor (*A^0^*) both in experiments started with *G^1^* (Fig. 4a) and *G^2^* (Fig. S2). Here, the ancestor lineages were, again, outcompeted by new lineages. However, in these experiments, the adapted lineages were not specialists that did very poorly in the opposing phenotype but were instead variations of generalists with higher fitness in either one or both phenotypes compared to the original ancestor *A^0^* (Fig. 4b, Table S1c) and the recent ancestor *G^1^* (Fig. 4c, Table S1d). As can be seen on the y-axis of Fig. 4c, strains *G^1^_G1_, G^1^_G2_, G^1^_G3_* and *G^1^_G4_* outcompeted *G^1^* in the swimming assay, but all except *G^1^_G4_* had a slightly decreased competitive fitness in the biofilm assay compared to *G^1^. G^1^_G4_* was more fit in the biofilm assay compared to *G^1^_G1_, G^1^_G2_*, and *G^1^_G3_* but was less fit in the swimming assay. Gauging the c-di-GMP levels relative to those of the original ancestor (*A^0^*) (Fig. 4c, Table S1e) showed that all derivatives of *G^1^* had a lower level of c-di-GMP compared to *G^1^* in the swimming assay.

**Figure 4.**
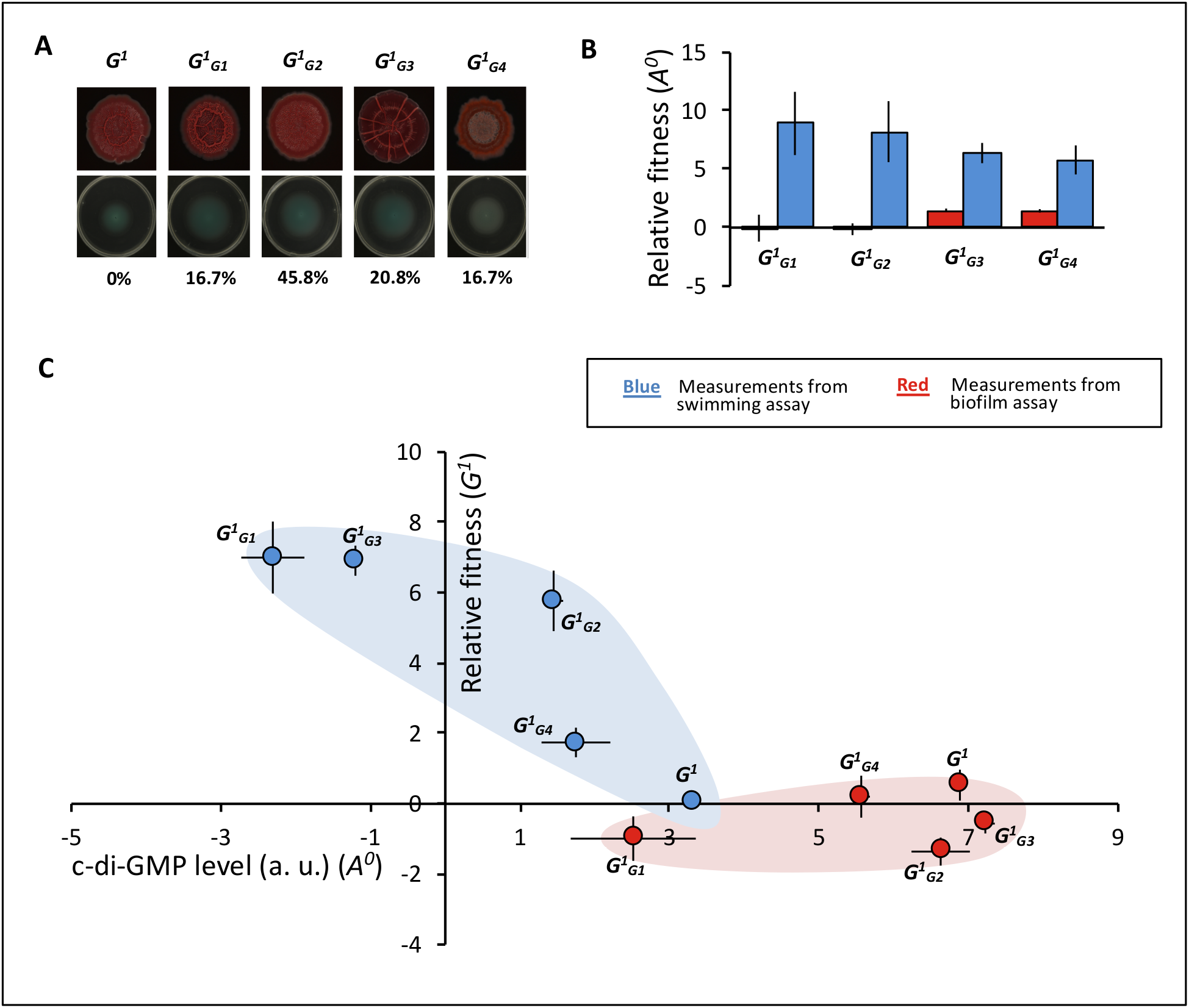
Resettlement of generalists with improved responsive phenotypic switching and subsequent exposure to changing environments selected for a heterogeneous population of generalists. **a**) Colony morphology and swimming phenotypes of derivatives of *G^1^*. Percentages indicate the proportion of the cell lineage after 11 passes. **b**) Competitive fitness *G^1^* and derivatives relative to the original ancestor (*A^0^*). **c**) The competitive fitness of isolates (n = 6), relative to their immediate ancestor *G^1^* as a function of c-di-GMP levels (n = 3), relative to *A^0^*. See Table S1c, S1d & S1e for *P*-values of oneway ANOVA with post hoc Dunnett test.

In the biofilm assay the two most abundant strains, *G^1^_G2_* and *G^1^_G3_*, had c-di-GMP levels comparable to *G^1^* while the levels of *G^1^_G1_* and *G^1^_G4_* were lower than *G^1^*, yet higher than *A^0^*. The above showed that after resettlement of *G^1^*, selection in changing environments favored generalist strains that adapted towards improved swimming motility while mostly retaining their biofilm phenotype. As seen in Fig. 4, among the derivatives of *G^1^*, levels of c-di-GMP were negatively correlated with the competitive fitness in the swimming assay (Fig. 4c, *Pearson’s r* **= 0.88,** *P* = 0.05). No correlation between the competitive fitness in the biofilm assay and c-di-GMP levels was found (Fig. 4c, **Pearson’s *r* = 0.29,** *P* = 0.635).

### Exposure to a third environment attests a tradeoff between maximizing fitness of fewer phenotypes versus averaging fitness of many phenotypes

In accordance with theory, our initial experiments show that a heterogeneous environment can select for individual organisms capable of expressing multiple phenotypes (generalist), but that the optimal fitness peak(s) of organisms with fewer phenotypes was higher comparatively. To test this relationship, we exposed the different strains to a third environment in the form of a murine burn wound model to which the evolved strains had not been exposed during the evolution experiments. *P. aeruginosa* PA14 (*A^0^*) was originally isolated from a burn wound and should be more fit in this third environment compared to the evolved strains. This is because the aforementioned theory predicts a trade-off for increased fitness in the swimming and/or biofilm assay should occur at the cost of optimal fitness through other phenotypes not selected for - a third environment.

We observed the highest mortality rate among mice infected with the ancestral PA14 (*A^0^*), followed by those infected with the swimming specialist (*S^3^*) (Fig. 5a). The groups infected with generalist (*G^1^*) or generalist (*G^2^*) had somewhat similar survival rates, whereas no mortality was observed in the group infected with the biofilm specialist (*B^2^*). In addition to virulence, we found that *A^0^* was more fit during burn infection compared to the other strains (Fig. 5b). Among the evolved strains, there was an interconnection between both CFUs and virulence, and the strains’ ability to swim. This was not the case for *A^0^* which was moderately good at swimming but highly virulent and fit.

**Figure 5.**
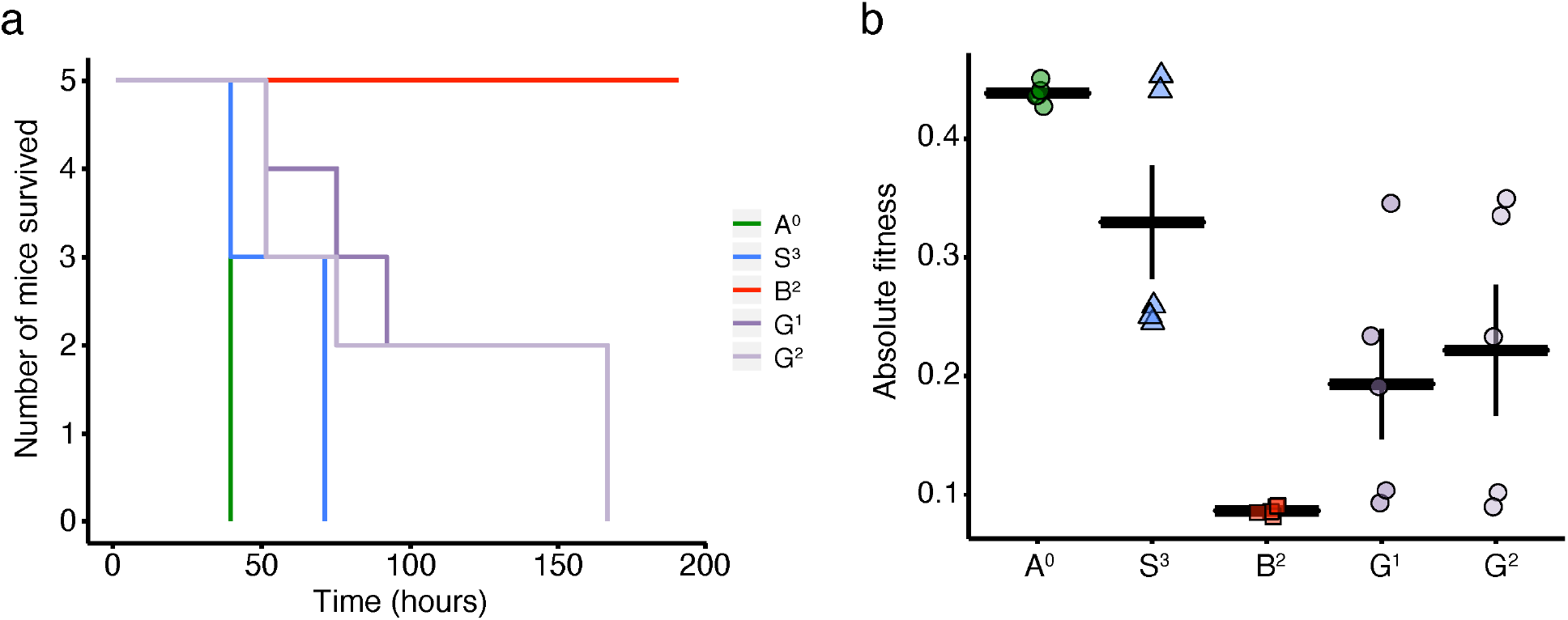
Consequences of specialization or generalization on virulence and fitness during burn wound infection. a) Survival curves of mice infected with ancestral PA14 (*A^0^*), swimming specialist (*S^3^*), biofilm specialist (*B^2^*), generalist (*G^1^*), or generalist (*G^2^*). b) Bacterial fitness indices in the mice shown in panel (a). Absolute fitness is defined as the LN(CFU_final_/CFU_initial_)/time, where CFU_final_ is the bacterial load per gram of tissue at time-of-death, CFU_initial_ is the bacterial load used to initiate the infection, and Time is the time-of-death (hours). Each data point represents values obtained from infection of an individual mouse. The CFU_final_ is the sum of the CFU/g of the three body sites sampled (spleen, tissue distal to the burn wound, and center of the burn wound). See Table S1f for adjusted *P*-values calculated using general linear hypotheses test with manual contrast, error bars denote ±SEM, n=5 for each group.

## Discussion

In accordance with theory on adaptation in changing environments^2^, our data indicate that during initial adaptation without spatial isolation, generalists will not be able to obtain as high a fitness peak in the two environments as the respective specialists. We find that opposing phenotypes that normally cannot be expressed simultaneously by one individual, such as the biofilm and swimming phenotypes^13,14^, enhance the likelihood of a “jack of all trades, master of none” predicament following results based on modeling^15^. During the initial experiments, we found that specialists outcompeted generalists during global competition despite that the generalists gained a higher average fitness compared to the ancestor in the two environments. Yet, adaptation to a higher level of responsive phenotypic switching, compared to the ancestor, occurred based on only a few mutations. These responsive generalists controlled their intracellular level of c-di-GMP more efficiently compared to the ancestor and the specialists. In accordance, further evolution of generalists after resettlement showed that the correlation between c-di-GMP levels and biofilm and swimming phenotype (Fig. 4) was retained among these strains. Exposing the evolved strains to a third environment showed that even the evolved generalists were subjected to a trade-off resulting in reduced virulence and fitness during infection (Fig. 5). This further strengthens the conclusion that a trade-off exists between maximizing fitness of fewer phenotypes versus averaging fitness of many phenotypes.

Mimicking relocation by restarting the changing environment evolution experiments using an adapted generalist showed that specialists that reduced the ability to perform opposing phenotypes were selected against. This may suggest that specialists were less likely to evolve based on the genetic background of the adapted generalists (*G^1^* and *G^2^*), but it is more likely that selection favored variations of generalists because the fitness peaks of the generalists in these experiments were closer to those possible to be obtained by the specialists. We speculate that there are a fewer numbers of mutations that can generate generalists compared to specialists, so the probability of generalists arising would have been lower than specialists, supporting the above reasoning. Resettlement is a hallmark of the generalists^16^ and facilitates spatial isolation. Therefore, within a limited number of generations after resettlement, a shift occurs from global to a more local competition against derivatives of the immediate ancestor^17^, and this may increase the probability of the successful evolution of generalists.

Bearing in mind that generalists during the experimental evolution were competing simultaneously with both biofilm and swimming specialists whose fitness was comparatively higher in the environment they are specialized to (Fig. 2) suggests that a prerequisite for the success of specialists is that the number of dedicated specialists is equal to the number of possible niches and that each specialist has a higher fitness peak in the relevant environment compared to a generalist. Importantly, this implies that the establishment of one type of specialist (e.g. biofilm) depended on the other type of specialist (e.g. swimming) to efficiently outcompete the generalists during environmental change.

Many environmental cues induce each of the two opposing phenotypes and the regulatory machinery that orchestrates the shift is complex. However, the second messenger, c-di-GMP, seems to be ubiquitously involved in facilitating this shift in the majority of bacteria, where high levels of c-di-GMP typically induce biofilm formation while low levels induce planktonic and motile states^13^. In *P. aeruginosa* the Wsp chemosensory system is important for mediating the shift between the two states by controlling the intracellular level of c-di-GMP^18^. The Wsp system consists of seven proteins: WspA-F and WspR. WspA is a membrane-bound methyl-accepting chemoreceptor that responds to less well-defined external signal(s)^19,20^. WspC is a methyltransferase, and WspF is a methylesterase that regulates the activity of WspA by altering its methylation state. When hypermethylated, WspA activates WspR by phosphorylation. WspR is a diguanylate cyclase that upon activation synthesizes c-di-GMP, inducing the biofilm state.

The shift between sessile and planktonic states are key to the fitness of bacteria and the complex regulatory systems that control this switch suggests that responsive phenotypic switching is a successful way of responding to environmental change as these sensing systems otherwise are too expensive to utilize and maintain in terms of energetics. Altering the levels of second messengers such as cyclic di-GMP has been argued to be an energetically relatively cheap way to obtain plasticity^21^. We suggest that this may be why mutations that influence the c-di-GMP output were associated with adaptation towards increased responsive switching^22^. While c-di-GMP is important in the regulation of the planktonic to biofilm switch, many other systems/genes are also known to affect these phenotypes. Here we followed the evolution of one population and can therefore not say if the observed adaptive trajectory is common for all *P. aeruginosa* in general or if mutations not associated with c-di-GMP system, may also generate generalists. Nonetheless, this study is to our knowledge the first to show empirically and in real-time, that generalists evolve specifically under heterogeneous conditions.

The *fleN* (FleN_175c>G_) mutation made *S^1^* a better swimmer compared to the ancestor (*A^0^*) due to multi-flagellation. Interestingly, if *fleN* is knocked out, cells become multi-flagellated but immobile^12^. Curiously, the c-di-GMP level of *S^1^* was lower compared to *A^0^* in the swimming assay but was similar in the biofilm assay where *S^1^* was outcompeted by *A^0^*. This suggests that for the strains with the *fleN* mutation to become better at forming a biofilm, the c-di-GMP level needed to be additionally enhanced in the biofilm assay to compensate for the reduction that followed the *fleN* mutation. Missense (and non-nonsense) mutations in the Wsp system seemed to be a primary target for this. Here, two outcomes were observed, either 1) mutations lead to a highly fit biofilm genotype that cannot reduce its c-di-GMP level sufficiently in the swimming assay and thus does poorly here, or 2) Wsp mutations that, despite facilitating high c-di-GMP levels in the biofilm assay, still allows the strain to reduce c-di-GMP levels sufficiently to also swim well. Data presented in Fig. 3 and 4c show that c-di-GMP levels of generalists and biofilm specialists in the biofilm assay reached approximately the same heights. However, in Fig. 3, swimming specialists had lower c-di-GMP levels in the swimming assay compared to generalists. This seemed to change after adaptation following resettlement, where the c-di-GMP levels of *G^1^* derived strains were lower. Further adaptive specialization as a swimming specialist also seemed to influence systems that affect c-di-GMP levels, at least this may explain the difference in fitness between *S^1^* and *S^2^*. It remains to be investigated how the mutation in the ribosomal binding site of *groES* furthered the swimming ability of the hyperflagellated *fleN* mutated genotype.

Infection experiments using a murine burn wound model (Fig. 5) showed that the ancestral PA14 (*A^0^*) was more virulent and fit in this environment compared to any of the experimentally evolved strains. Virulence is, like motility and biofilm formation, regulated by c-di-GMP and we, therefore, speculate that the ancestor *A^0^* was more virulent and fit during infection because it expressed apt phenotypes more optimally, by achieving very specific and not aberrant c-di-GMP levels when compared to the evolved strains.

## Materials and Methods Evolution experiments

*Pseudomonas aeruginosa* UCBPP-PA14 (*A^0^*) was used as the ancestral strain (*A^0^*) for evolution experiments other than the resettlement experiments where generalists *G^1^* and *G^2^* were used as the ancestral strain. All experimental evolutions were performed in two independent replicates. Overnight cultures of ancestral strains were diluted to an OD_600_ of 0.15 and inoculated into the swimming and pellicle assay as described below. Experimental evolutions in changing environments were started with the swimming assay and screening of the colony morphology and swimming motility were done after growth in the pellicle assay. Screenings were performed after 11, 23 and 35 passes in experiments started with *A^0^*, while screenings were done after 11 passes in experiments started with *G^1^* and *G^2^*. Control experiments that mimicked stable environments were performed using *A^0^* as the original inoculum and cultures were passed from and to the same assay. This was done in shaken cultures, the pellicle assay, and the swimming assay. 35 passes were done for the control experiments. When screening colony morphology and swimming motility between 30 and 60 random colonies were isolated and re-streaked a minimum of two times.

### Colony morphology assay

Ten microliters of overnight cultures (250rpm, 37 °C) adjusted to OD_600_ 0.15 were spotted in the center of agar plates (94 x 16 mm) with 40 mL colony morphology assay medium; 10 g/L tryptone (Technova), 10 g/L agar (Technova), 40 ug/mL Congo red (CR), and 20 ug/mL Coomassie brilliant blue (CB). Colonies were grown at 24 °C for up to 5 days.

### Pellicle assays

Two microliters of overnight cultures (250rpm, 37°C) adjusted to OD_600_ 0.15 were inoculated into 5 mL of Miller LB (Microbiology) broth. Tubes were left unshaken at 37°C for 24h. After incubation, the tube was vortexed vigorously and homogenized by pipette mixing before either being passed or processed by flow cytometry.

Competitive fitness in pellicle assays: when competing two strains a *gfp*- and a *mCherry-tagged* version of both strains was used; strain A_gfp_ was competed against B_mCherry_ and A_mcherry_ against Bgfp. Overnight cultures (250rpm, 37 °C) of all 4 strains were adjusted to OD_600_ = 0.15. The two sets of strains were mixed in 1:1 ratios and inoculated into the pellicle assay. Start ratios were measured and recorded by flow cytometry. After incubation, the ratio was again recorded by flow cytometry. CFUs were obtained for control experiments and the number of *gfp* and *mCherry* tagged bacteria were calculated based on identifying *gfp*-tagged CFUs using a Transillumination DR-88M.

### Swimming motility assays

Two microliters of overnight cultures (250rpm, 37 °C) adjusted to OD_600_ 0.15 were stabbed into the center of agar plates (94 x 16 mm) with 25 mL swimming agar; 5 g/L Miller LB broth (Microbiology), and 3 g/L agar (Technova). Agar plates were left at 37 °C for 16h. After incubation cells imbedded in the soft agar were transferred to sterile 50 mL Falcon tubes by use of a sterile spoon and homogenized by vortex mixing before either being passed or processed by flow cytometry.

Competitive fitness in swimming assays: when competing two strains a *gfp*- and a *mCherry-tagged* version of both strains was used; strain Agfp was competed against B_mCherry_ and A_mCherry_ against Bgfp. Overnight cultures (250rpm, 37 °C) of all 4 strains were adjusted to OD_600_ = 0.15. The two sets of strains were mixed in 1:1 ratios and inoculated into the swimming assay. Start ratios were measured and recorded by flow cytometry. After incubation, the ratio was again recorded by flow cytometry. CFUs were obtained for control experiments and the numbers of *gfp* and *mCherry* tagged bacteria were calculated based on identifying *gfp-tagged* CFUs using a Transillumination DR-88M.

### Burn wound infection murine model

The murine model of burn wound infection was adapted from the model described by Stieritz and Holder^23–25^. Burn experiments were conducted with adult female Swiss Webster mice (Charles River Laboratories, Inc.) weighing 20-25 g. Mice were anesthetized by intraperitoneal injection of 5% sodiumpentobarbital (Nembutal; Oak Pharmaceuticals, Inc.) at 5 mg/mL before their backs were shaved, and the hair was cleanly removed with a depilatory agent. As a preemptive analgesic, 0.05 mL of topical anesthetic (bupivacaine [0.25%] with lidocaine [2%]) was applied to the area to be wounded. Thermal injury was induced by placing an exposed area of the shaved skin (approximately 15% total body surface area) in a 90°C water bath for 10 s. This scald injury is nonlethal but induces a third-degree (full-thickness) burn. Immediately, mice were given a subcutaneous injection of sterile 0.5 mL 0.9% NaCl saline solution on a distal site anterior of the burn wound.

*P. aeruginosa* inocula for challenge in thermally injured mice were prepared as previously described^24,25^. Mice were infected with 10^2^ CFU of *P. aeruginosa* in 100 μL 1× PBS of ancestral PA14 (*A^0^*), swimming specialist (*S^3^*), biofilm specialist (*B^2^*), generalist (*G^1^*), or generalist (*G^2^*). Mice were monitored for 192 hours (8 days) post-injury and infection. At the specified end time point or in the event mice became moribund, mice were euthanized by intracardiac injection of 200 μL (390 mg/mL) Fatal-Plus sodium-pentobarbital (Vortech Pharmaceuticals, Ltd.). Organs were gross examined at the time-of-death, the spleen, tissue distal to the burn wound, and center of the burn wound were excised, homogenized and assessed for bacterial load per gram of tissue by enumeration of colony forming units. The experiment was performed twice with five mice per group to ensure reproducibility. Animals were treated humanely and in accordance with protocol 96020 approved by the Institutional Animal Care and Use Committee at Texas Tech University Health Sciences Center in Lubbock, Texas, USA.

### Insertion of *gfp* and *mCherry* into the chromosome of *P. aeruginosa* strains

The *gfp* or *mCherry* was inserted into the chromosome of *P. aeruginosa* strains by using a mini-CTX2 based system^26^. More detailed procedures can be found in Almblad et al.^26^ *P. aeruginosa* strains were made electrocompetent by washing (2 min. at 10000 *g*) 5 mL overnight cultures 3 times in 300nM glycerol. Cells were then pelleted, the supernatant removed, and resuspended in 50 μL 300nM glycerol. Next, plasmid mini-CTX2T2.1-GW::Ptrc-*GFP* or mini-CTX2T2.1-GW::Ptrc-*mCherτy* was transformed into the strains. Transconjugants were selected for on LB agar supplemented with 100 ug/mL tetracycline. Hereafter pFLP2 was used to remove the tetracycline resistance cassette following procedures described by^27^, producing *gfp* or *mCherry* labeled, but antibiotic-sensitive *P. aeruginosa* strains.

### Genome sequencing and identification of mutations

DNA was extracted from overnight cultures of *P. aeruginosa* strains using the Qiagen DNeasy Blood & Tissue kit (Qiagen, cat. no. 69504). The whole-genome sequencing libraries were prepared using the Nextera XT DNA library preparation kit (Illumina, USA) and subsequently quantified by Fragment Analyzer™ (Advanced Analytical Technologies Co.). Sequencing was done as 2 x 250-base paired-end reads using the Illumina MiSeq platform (Illumina). Standard protocols were used for all of the above kits as provided by the manufacturer. Data were analyzed using CLC Genomic Workbench V7.5.1 (CLCBio). Obtained reads were trimmed and normalized. Briefly, reads were trimmed removing adapter sequences and discarding those of low quality using “trim sequences” tool (settings: ambiguous limit = 2, quality limit = 0.05). First, trimmed paired and orphan reads of the ancestor *A^0^*, PA14 WT were mapped to the genome sequence of *Pseudomonas aeruginosa* UCBPP-PA14 (Accession number: NC_008463) using the “Map Reads to Reference” tool (default settings). Next, the same was done for trimmed paired and orphan reads of strains; *S^1^*, *S^2^, S^3^, B^1^, B^2^, G^1^*, & *G^2^*. The lowest percentage of reads that matched was 99.9% (Table S3). Following this, a re-sequencing analysis was performed using the “basic variance detection” tool. The “probabilistic variant detection” tool (min. coverage 10 bp, variant probability 0.95) was then used to identify variations in the genomes of strains *S^1^, S^2^, S^3^, B^1^, B^2^, G^1^*, & *G^2^* relative to the ancestral strain. The variance was verified or rejected by manually inspecting the individual mapped sequences. In addition, the trimmed paired and orphan reads were also assembled *de novo* using the “de novo assembly” tool (Min. contig length 500bp, Map reads back to contig; default settings). Reads were mapped back to contigs (min. length of contigs was set to 500 bp otherwise default settings). The *de novo* sequences were then inspected, and mutations verified.

### Cyclic di-GMP reporter constructs

Reporter pCdrA::*gfp*(ASV)^S^, a fluorescence-based reporter for gauging cyclic di-GMP levels^28^, was introduced into the *P. aeruginosa* strains via electroporation and selected for on LB agar supplemented with 100 ug/mL gentamicin. Strains were made electrocompetent as described above.

### Flow cytometry

Flow cytometry was performed on a BD FACSAria IIIu (Becton Dickinson Biosciences, San Jose, CA, USA). Settings and voltages; forward scatter = 505V, side scatter = 308V, green fluorescence (bandpass filter 530/30 nm) = 508V, and red fluorescence (bandpass filter 610/20 nm) = 500V. A 70 um nozzle was used at a sheath fluid pressure of 70 psi. The BD FACSDiva software v.6.1.3 was used for operating and analyzing results.

Competition assays: gates were defined using *gfp* and *mCherry* tagged and non-tagged versions of the *P. aeruginosa* PA14 wild-type (*A^0^*). When obtaining ratios of green and red fluorescent *P. aeruginosa*, only fluorescent events were counted and they were only counted if the events were not both green and red. Gates and quantifications were verified by CFU counting.

Cyclic di-GMP measurements: a *gfp* threshold was defined using a non-tagged version of *P. aeruginosa* UCBPP-PA14 (*A^0^*) and the same strain carrying the pCdrA::*gfp*(ASV)^S^ reporter. Cells from shaken cultures, the pellicle of static cultures, and the swimming assay were used when setting the threshold. Samples were taken at different distances from the edge of the advancing swimming zone down to the center of the swimming zone. The pCdrA::*gfp*(ASV)^S^ reporter is based on the expression of an unstable version of GFP and the number of green events that exceeded the threshold was used as a measure of population-averaged c-di-GMP levels.

### Calculating relative competitive fitness and relative community c-di-GMP levels

The relative competitive fitness of strains was calculated as ratios of *gfp* and *mCherry* events (number of mutants divided by the number of ancestors) when biofilm and swimming assays were incubated (start ratio) and after incubation (end ratio) and this number was then log2 transformed. Relative competitive fitness = log2[(start ratio)/(end ratio)]. The relative community c-di-GMP levels were calculated by log2 transforming the number of *gfp* events of the derived strain divided by those of the ancestor (always the *A^0^* strain).

### Imaging using transmission electron microscopy

Transmission electron microscopy was performed directly from overnight cultures. 400 mesh carbon-coated Cu/Rh grids were glow discharged for 30 s immediately prior to use. Grids were incubated on 10 μL of overnight culture for 1 minute before blotting the liquid media off with filter paper. Subsequently, each grid was incubated in 10 μL of 1% uranyl acetate for 2 min before blotting. Finally, each grid was blotted with 10 μL of ddH2O, and blotted dry with filter paper. Grids were left to dry before image acquisition. Images were acquired using a CM 100 transmission electron microscope and captured using an Olympus Veleta camera. All acquired photos were used to assess the distribution of the number of flagella by manual counts.

### Statistical analyses

Data and statistical analyses were performed using R version 3.5.029^29^. Statistical tests used for the analysis of data are identified in the legend of each figure. Differences of *P*<0.05 were considered significant.

## Acknowledgements

We would like to thank Dr. Henrik Almblad and Prof. Tim Tolker-Nielsen very much for sharing their constructed vectors CTX2T2.1-GW::Ptrc-*GFP*, mini-CTX2T2.1-GW::Ptrc-*mCherry* and pCdrA::*gfp*(ASV)^S^. Also, a big thanks to Anette Hørdum Løth and Luma George Odish for excellent help with laboratory experiments. This work was supported by research grants from the Danish Council for Independent Research (DFF-1323–00235, SIMICOM); Lundbeckfonden (SHARE, R250-2017-1392); and VILLUM FONDEN (00028107).

