## Supplement information for "Adaptive evolution of phenotypic switching during environmental change"

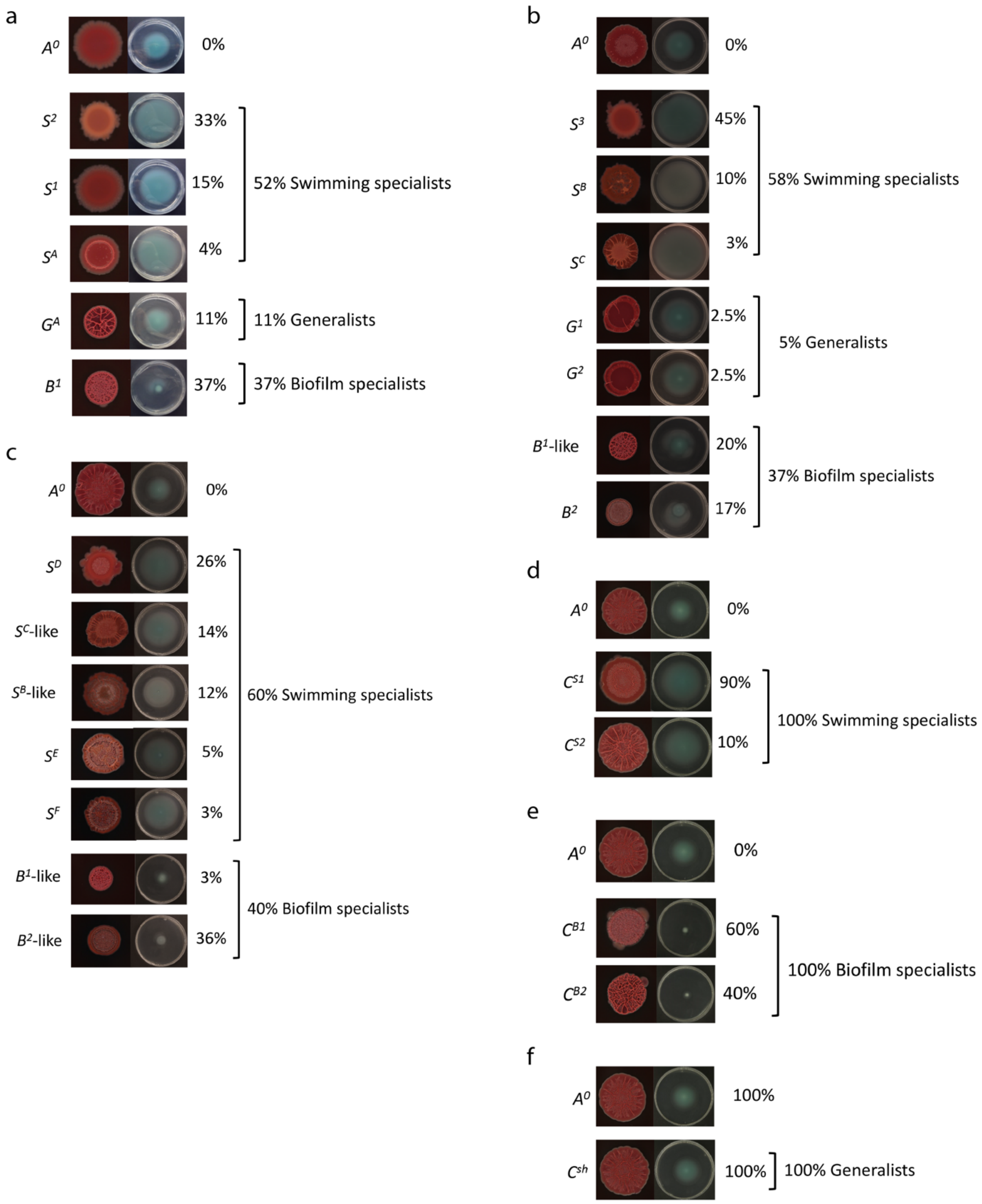


**Figure S1. Proportion of cell lineages after the respective number of passes. a)** After 11 passes in changing environments. **b)** After 23 passes in changing environments. **c)** After 35 passes in changing environments. **d)** After 11 passes in swimming assays. **e)** After 12 passes in pellicle assays. **f)** After 23 passes in shaken cultures.

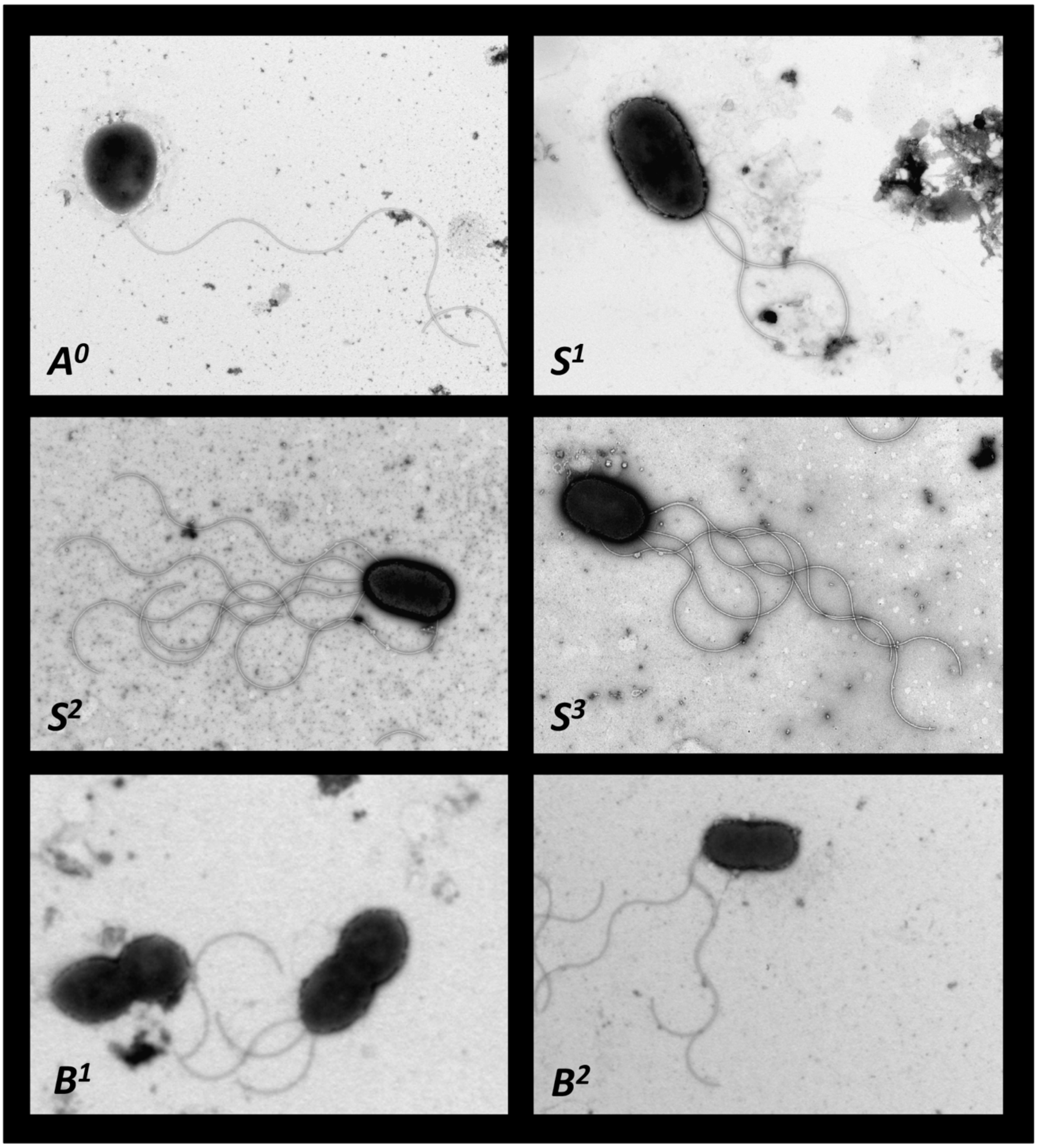


**Figure S2. Transmission electron microscopy (TEM) pictures of select strains.**

### Table S1a. Comparison of relative fitness values of mutants and $A^0$

| Strain | Swimming assay | Biofilm assay |
|---|---|---|
| $G^1$ | $P = 0.0254$* | $P < 0.001$*** |
| $G^2$ | $P = 0.0477$* | $P = 0.001$** |
| $S^1$ | $P < 0.001$*** | $P = 0.089$· |
| $S^2$ | $P < 0.001$*** | $P < 0.001$*** |
| $S^3$ | $P < 0.001$*** | $P < 0.001$*** |
| $B^1$ | $P < 0.001$*** | $P < 0.001$*** |
| $B^2$ | $P < 0.001$*** | $P < 0.001$*** |

One-way ANOVA with post hoc Dunnett test, $n = 6$.

### Table S1b. Comparison of relative c-di-GMP levels of mutants and $A^0$

| Strain | Swimming assay | Biofilm assay |
|---|---|---|
| $G^1$ | $P = 0.773$ | $P < 0.001$*** |
| $G^2$ | $P = 0.518$ | $P < 0.001$*** |
| $S^1$ | $P < 0.001$*** | $P = 0.376$ |
| $S^2$ | $P < 0.001$*** | $P < 0.999$ |
| $S^3$ | $P = 0.001$** | $P < 1.0$ |
| $B^1$ | $P < 0.001$*** | $P < 0.001$*** |
| $B^2$ | $P = 0.004$** | $P < 0.001$*** |

One-way ANOVA with post hoc Dunnett test, $n = 4$.

### Table S1b. Comparison of relative fitness values of mutants and $A^0$

| Strain | Swimming assay | Biofilm assay |
|---|---|---|
| $G1^{G1}$ | $P < 0.001$*** | $P = 0.981$ |
| $G1^{G2}$ | $P < 0.001$*** | $P = 0.895$ |
| $G1^{G3}$ | $P < 0.001$*** | $P = 0.001$** |
| $G1^{G4}$ | $P < 0.001$*** | $P < 0.002$** |

One-way ANOVA with post hoc Dunnett test, $n = 6$.

### Table S1d. Comparison of relative fitness values of mutants and $G^1$

| Strain | Swimming assay | Biofilm assay |
|---|---|---|
| $G1^{G1}$ | $P < 0.001$*** | $P < 0.001$*** |
| $G1^{G2}$ | $P < 0.001$*** | $P < 0.001$*** |
| $G1^{G3}$ | $P < 0.001$*** | $P = 0.002$** |
| $G1^{G4}$ | $P < 0.001$*** | $P < 0.602$ |

One-way ANOVA with post hoc Dunnett test, *n* = 6.

#### Table S1e. Comparison of relative c-di-GMP levels of mutants and $A^0$

| Strain | Swimming assay | Biofilm assay |
|---|---|---|
| $G1^{G1}$ | $P < 0.001$*** | $P < 0.001$*** |
| $G1^{G2}$ | $P < 0.001$*** | $P < 0.001$*** |
| $G1^{G3}$ | $P = 0.003$** | $P < 0.001$*** |
| $G1^{G4}$ | $P < 0.001$*** | $P < 0.001$*** |

One-way ANOVA with post hoc Dunnett test, *n* = 3.

#### Table S1f. Comparison of absolute fitness of mutants and $A^0$

| Comparison | Burn infection |
|---|---|
| $A^0$-$S^3$ | $P = 0.1823$ |
| $A^0$-$B^2$ | $P < 0.001$*** |
| $A^0$-$G^1$ | $P < 0.001$*** |
| $A^0$-$G^2$ | $P = 0.0029$ |

Adjusted *P*-values calculated using general linear hypotheses test with manual contrast, n=5.

#### Table S2. Identified mutations based on full genome sequencing

| Strain | Pass # | Mutation | Affected protein | Mutation nt | Mutation aa | Domain |
|---|---|---|---|---|---|---|
| $G^1$ | 23 | $SNP_1$ | FleN (PA14_45640) | 4059726 A>C | 175 C>G | cd02038 |
| | | $SNP_2$ | WspA (PA14_16430) | 1406990 T>G | 449 F>C | cd11386 |
| $G^2$ | 23 | $SNP_1$ | FleN (PA14_45640) | 4059726 A>C | 175 C>G | cd02038 |
| | | $SNP_3$ | WspF (PA14_16480) | 1412596 A>G | 185 Q>R | Pfam 01339 |
| $S^1$ | 11 | $SNP_1$ | FleN (PA14_45640) | 4059726 A>C | 175 C>G | cd02038 |
| $S^2$ | 11 | $SNP_1$ | FleN (PA14_45640) | 4059726 A>C | 175 C>G | cd02038 |
| | | $SNP_4$ | RBS of *groES* (PA14_57020) | 5081023 A>C | - | - |
| $S^3$ | 23 | $SNP_1$ | FleN (PA14_45640) | 4059726 A>C | 175 C>G | cd02038 |
| | | $SNP_4$ | RBS of *groES* (PA14_57020) | 5081023 A>C | - | - |
| | | $DEL_1$ | PA14_29800 | 2580934 ΔCGAGCTGGG | 262 ΔRAG | cd06225 |
| | | $SNP_5$ | PA14_32850 | 2875326 | Missense | - |
| $B^1$ | 11 | $SNP_1$ | FleN (PA14_45640) | 4059726 A>C | 175 C>G | cd02038 |
| | | $SNP_6$ | WspA (PA14_16430) | 1406789 T>C | 382 V>A | cd11386 |
| $B^2$ | 23 | $SNP_1$ | FleN (PA14_45640) | 4059726 A>C | 175 C>G | cd02038 |
| | | $SNP_6$ | WspA (PA14_16430) | 1406789 T>C | 382 V>A | cd11386 |
| | | $SNP_7$ | PA14_45960 | 4085368 C>T | 231A>T | cd02038 |

Colors highlight similar mutations in different strains.
nt: nucleotide, aa: amino acid, SNP: single nucleotide polymorphism, DEL: deletion.

### Table S3. Mapping summary report

| Strain | Total reads after trimming | Number of reads match | Average coverage |
|---|---|---|---|
| $A^0$ | 1271202 | 1269398 | x52 |
| $G^1$ | 2518088 | 2515557 | x70 |
| $G^2$ | 3810762 | 3807083 | x103 |
| $S^1$ | 5180430 | 5174869 | x140 |
| $S^2$ | 3572136 | 3568627 | x116 |
| $S^3$ | 4762343 | 4758130 | x133 |
| $B^1$ | 3552979 | 3549348 | x96 |
| $B^2$ | 3010215 | 3007620 | x83 |

Reads were mapped against the complete genome of *Pseudomonas aeruginosa* UCBPP-PA14 (Accession number: NC_008463)
